# Use of SNPs in a low diversity system for genetic monitoring and identifying a successful translocation event in the Paiute Cutthroat Trout (*O. henshawi seleniris*)

**DOI:** 10.64898/2026.01.07.697447

**Authors:** Yingxin Su, Melanie E.F. LaCava, Matthew A. Campbell, Chad Mellison, Robert G. Titus, Jeff Rodzen, Andrea D. Schreier, Amanda J. Finger

## Abstract

**Objective:** Maintaining genetic diversity in small, isolated populations is a key goal in conservation genetics. Genetic monitoring can guide management decisions, but comparisons across studies and time points are often complicated by differences in genetic markers and data sets. We aimed to develop a consistent, replicable set of genetic markers to support long-term monitoring of Paiute Cutthroat Trout (*Oncorhynchus henshawi seleniris*), a federally threatened subspecies persisting in a network of isolated refuge populations in California, USA.

**Methods:** We used RAD (Restriction site-associated DNA) sequencing to identify single nucleotide polymorphisms (SNPs) from 476 individuals representing eight extant refuge populations. Filtering steps included the removal of paralogous loci and exclusion of *F_ST_* outliers associated with sequencing batch effects. We selected a panel of 1,114 SNPs that were shared across all populations and validated the panel *in silico* by comparing population structure and genetic diversity estimates to those derived from the full RAD sequencing dataset.

**Results:** The candidate panel SNPs captured key patterns of genetic differentiation and diversity consistent with previous studies and the larger RAD dataset. We present updated baseline genetic metrics for each refuge population, providing a consistent reference point for future monitoring efforts. We also demonstrate the ability of our candidate panel SNPs to detect successful spawning after a translocation event.

**Conclusion:** This study provides a set of SNPs tailored for monitoring genetic diversity and structure in Paiute Cutthroat Trout refuge populations. Future work should include development of a high-throughput genotyping assay to implement these candidate panel SNPs for ongoing management. This would enable consistency over time with genetic studies and be a foundational tool for repeated evaluation of conservation actions such as reintroductions and augmentations.

**Lay Summary:** We identified genetic markers to help track and protect one of the rarest trout in the United States. This work supports long-term conservation by making it easier to monitor isolated populations and measure the success of efforts to restore the species.

## Introduction

Maintaining genetic diversity is a central goal when conserving threatened species, yet many of these species persist only in a handful of isolated “refuge” populations (Frankham 2005; Novak et al. 2020). Refuge populations can be *in-* or *ex-situ* and are typically established as both safeguards against extinction and reservoirs for recovery efforts (Meretsky et al. 2006; Todesco et al. 2016; Love Stowell et al. 2017). Despite their importance, refuge populations are vulnerable to genetic risks stemming from founder effects (Templeton et al. 2001; Fraser 2008) and small population sizes (Frankham 2005), which intensify the effects of genetic drift, eroding genetic diversity over time (Montgomery et al. 2000; Abascal et al. 2016). Furthermore, genetic drift drives genetic differentiation among small refuge populations over time such that what was once a large single population becomes subdivided into many small populations diverging away from each other. To mitigate these risks and ensure the long-term viability of refuge populations, genetic data should play a role in monitoring and management, guiding decisions about genetic rescue, augmentation, and recovery planning (Meffe and Vrijenhoek 1988; Hoffman et al. 2020; Cook et al. 2023).

The Paiute Cutthroat Trout (*Oncorhynchus henshawi seleniris*, hereafter PCT; Snyder 1933; Page et al. 2023) is a federally threatened subspecies of Lahontan Cutthroat Trout (*O. henshawi*) that exemplifies the challenges of managing refuge populations. PCT have a complex translocation history and limited genetic diversity, underscoring the importance of incorporating genetics into conservation and monitoring efforts. Historically, PCT had the narrowest range of any cutthroat trout, limited to a single drainage in the Carson River basin in the Sierra Nevada of east-central California (Figure 1). Their native distribution spanned less than 20 km of Silver King Creek, between the 25-foot Llewellyn Falls and a series of barriers in Silver King Canyon, known commonly as the Gorge (Behnke 2002; Figure 1B). The long isolation of PCT from other cutthroat trout (0.26-0.85 MYA; Saglam et al. 2017) likely led to its unique phenotype amongst Pacific trouts of an almost complete lack of body spotting and retention of parr marks (Snyder 1933; Behnke 1992). While PCT benefited from a remote location, repeated introductions of nonnative salmonids such as Rainbow Trout (*O. mykiss*) and Lahontan Cutthroat Trout (*O. henshawi*), throughout much of the 1900s threatened PCT with competition and hybridization (Schroeter 1998; USFWS 2004). Fortuitously, in 1912 PCT were introduced into headwater tributaries above barriers in the Silver King Creek watershed, which protected them by isolating them from nonnative salmonid pressure in lower reaches (Ryan and Nicola 1976). It is likely that by the 1920s and certainly by the time Snyder described the taxon in 1933, PCT no longer existed in their native habitat (Snyder 1933, Ryan and Nicola 1976) and were listed in 1967 as endangered under the federal Endangered Species Preservation Act of 1966 (USFWS 1967). In 1975 the species was down-listed under the Endangered Species Act of 1973 to threatened to allow management actions such as rotenone treatments to remove nonnative species, translocations, and the establishment of refuge populations (USFWS 1975). After a complex series of introductions and translocations over the last 100 years (details in Cordes et al. 2004; Finger et al. 2013; USFWS 2020), at the time of this study PCT persisted in nine refuge populations, five located in-basin (within the native Silver King Creek watershed) and four out-of-basin (Figure 1). Though the populations are genetically verified as unhybridized, earlier work showed that these populations are now differentiated from each other and have low levels of diversity (Cordes et al. 2004; Finger et al. 2013). Long-term viability of these individual, isolated refuge populations will always be threatened due to their small size and isolation (Lande 1998; Hedrick et al. 2000; Frankham 2005). With the understanding that genetic data are an important component of managing refuge populations, a 2013 Genetic Management Plan for the PCT used microsatellites to establish patterns of genetic structure and diversity among refuge populations and made suggestions for adaptive management strategies, including a recommendation to develop a single nucleotide polymorphism (SNP) panel for ongoing genetic monitoring (Finger et al. 2013).

**Figure 1.**
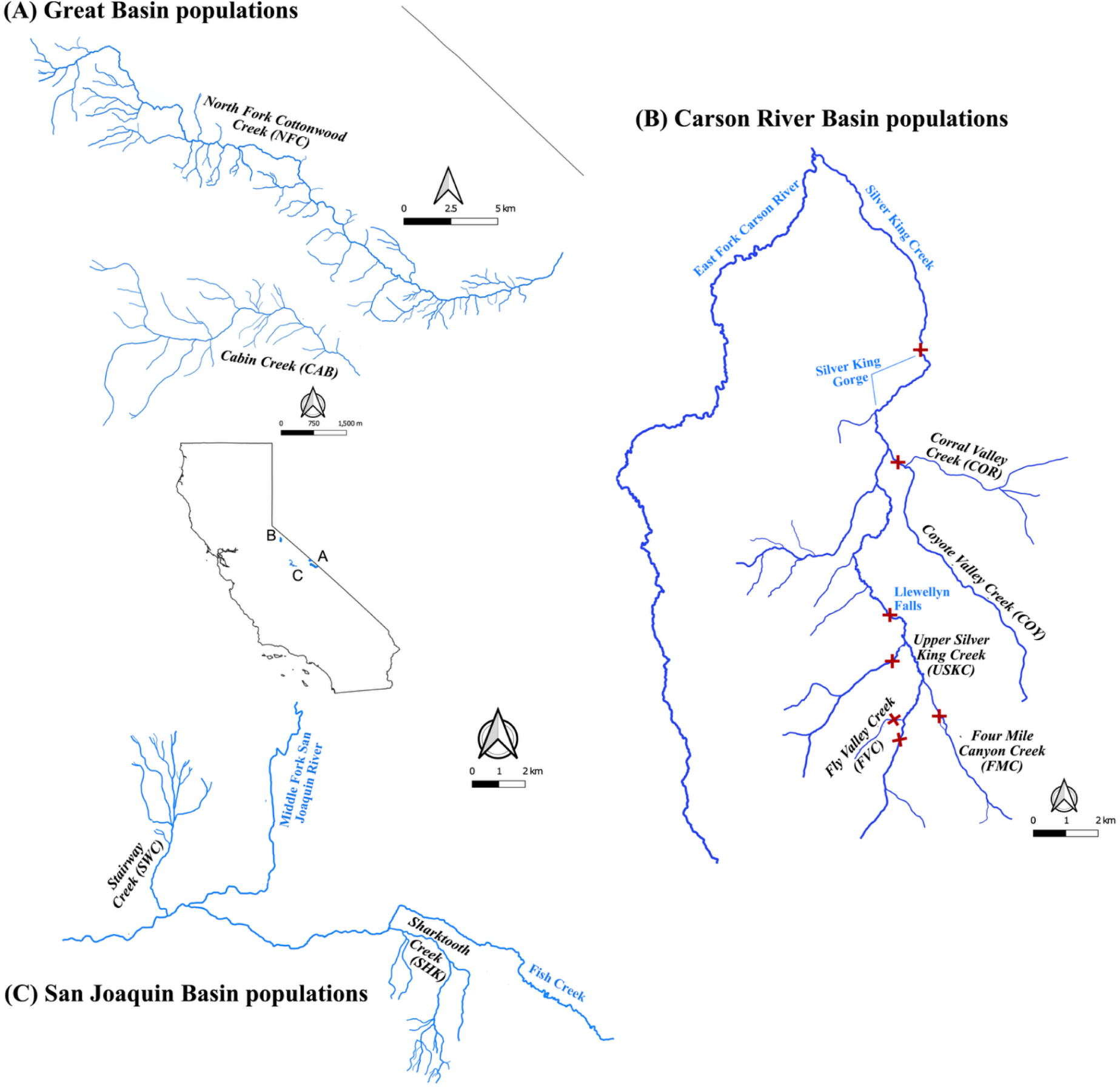
Geographic locations of PCT refuge populations (labeled in black) within three basins in the state of California, USA. (A) Two out-of-basin populations are located in the Great Basin, (B) five within-basin populations are located in the Carson River Basin, and (C) two out-of-basin populations are located in the San Joaquin Basin. Fish barriers are denoted by red hatch marks.

In an age of rapidly advancing sequencing technologies, numerous methods are available to develop SNP markers for genetic diversity monitoring of refuge populations. Although large, genome-wide datasets provide significant power and can be used without a reference genome, they often take significant time to produce, require substantial experience to analyze, and can produce inconsistent results making combining data sets difficult (Lowry et al. 2016; Flanagan and Jones 2018; O’Leary et al. 2018). Reduced representation sequencing methods such as RAD sequencing allow for more rapid and low cost identification of SNPs relative to whole genome sequencing, but it is less efficient for genetic monitoring, which may involve high-throughput genotyping of many individuals across multiple time points (Leigh et al. 2018). A well-designed panel of single nucleotide polymorphisms (SNPs) derived from RAD sequencing can provide a useful alternative for genetic monitoring that enables rapid genotyping with minimal loss of the power and accuracy associated with larger data sets (Meek and Larson 2019; Bootsma et al. 2020). For example, Mamoozadeh et al. (2023) recently surveyed genetic diversity across the range of Brook Trout (*Salvelinus fontinalis*) and produced a SNP dataset that could be leveraged to design SNP panels tailored for various conservation and management applications, illustrating the broad utility of this approach.

Though SNP panels can be a useful tool for routine monitoring, developing these panels can be challenging. For example, in cases where multiple small refuge populations have undergone significant drift due to isolation and low effective population sizes, the resulting low levels of genetic diversity may reduce the number of shared polymorphic sites among populations (Fraser 2008). In addition, evolutionary processes can introduce complexities into the genome that further complicate the accurate detection of informative SNPs. For example, in the family Salmonidae, an ancient whole genome duplication resulted in pervasive paralogous loci and residual tetrasomy in parts of the salmonid genome (Ferris and Whitt 1980; Macqueen and Johnston 2014; Campbell et al. 2019). As a result, special attention is required while developing a SNP panel to avoid collapsing paralogous loci, which would lead to inflated heterozygosity estimates (McKinney et al. 2017; Willis et al. 2017; O’Leary et al. 2018). Ideally, SNPs selected for genetic monitoring are unlinked, non-paralogous, present in every population of interest in a study system, and can be reliably genotyped. These characteristics enable unbiased estimates of genetic diversity, effective population size, population assignment, and genetic population structure (Bootsma et al. 2020; May et al. 2020).

In this study, we generated RAD sequencing data from eight extant refuge populations to identify a set of SNPs that, when genotyped with a genotyping-by-sequencing tool such as GT-seq (Campbell et al. 2015), are appropriate for genetically monitoring PCT. We validated the candidate panel *in silico* by comparing genetic structure and diversity metrics produced by the panel to past findings and to our full RAD sequencing dataset. We used the candidate panel SNPs to create an updated baseline for genetic structure and diversity, against which future genetic monitoring data can be compared since the previous baseline was based on 11 microsatellite loci. Finally, we used a 2017 translocation event to test the ability of our markers to detect whether translocated fish spawned in their new population. This study provides a critical tool for managers to monitor PCT refuge populations.

## Methods

### Sample collection, DNA sequencing, and initial quality filtering

Biologists from the California Department of Fish and Wildlife, U.S. Forest Service, and U.S. Fish and Wildlife Service collected fin clips from eight PCT refuge populations from 2017 to 2021 (Figure 1, Table 1). Four study populations are located within the Carson River Basin, including Silver King Creek above Llewellyn Falls (i.e., upper Silver King Creek), Fly Valley Creek, Corral Valley Creek, and Coyote Valley Creek (hereafter, we refer to these populations as ‘within-basin’). The remaining four populations are Sharktooth Creek and Stairway Creek, which are located in the San Joaquin River Basin, and North Fork Cottonwood Creek and Cabin Creek, which are located in the Great Basin (hereafter we refer to these four populations as ‘out-of-basin’). We excluded a ninth refuge population, the within-basin Four Mile Canyon Creek, from our study because we were unable to collect sufficient samples. An additional out-of-basin population, Delaney Creek, located within Yosemite National Park, Tuolumne County, California is presumed to be extinct due to competition with Brook Trout (USFWS 2004) and was not sampled. The USFWS (2004) document provides additional detail on the history and status of PCT populations beyond the scope of the current study.

**Table 1.** Refuge population sampling information including habitat length in kilometers (km), sample collection year, the number of samples sequenced, and the number of samples included in analyses after all bioinformatic filtering. The mean and variance of individual heterozygosity (het.) for each population was calculated using 1,114 panel SNPs. Some years were combined with consecutively sampled years due to low sample sizes. For additional context, we included observed heterozygosity (H_O_) based on microsatellite data from samples collected in 2010–2012 (Finger et al. 2013).

| Type | Basin | Population | Habitat length (km)<sup>1, 2</sup> | Sample collection year | Sample size | Sample size after filtering | Mean het. | Variance het. | H<sub>o</sub> from Finger et al. 2013 |
| --- | --- | --- | --- | --- | --- | --- | --- | --- | --- |
| Within -Basin | Carson River Basin | Corral Valley Creek (COR) | 3.6 | 2019 | 15 | 10 | 0.23 | 0.0020 | 0.41 |
|  |  |  |  | 2020 | 44 | 39 |  |  |  |
|  |  | Coyote Valley Creek (COY) | 4.9 | 2019 | 61 | 55 | 0.23 | 0.0025 | 0.46 |
|  |  | Fly Valley Creek (FVC) | 2.0 | 2019 | 10 | 9 | 0.23 | 0.0010 | 0.40 |
|  |  | Upper Silver King Creek (USKC) | 4.3 | 2019 | 24 | 20 | 0.27 | 0.0001 | 0.53 |
|  |  |  |  | 2021 | 116 | 87 | 0.27 | 0.0023 |  |
| Out-of -Basin | Great Basin | Cabin Creek (CAB) | 2.4 | 2020 | 8 | 8 | 0.24 | 0.0035 | 0.57 |
|  |  | North Fork Cottonwood Creek (NFC) | 5.5 | 2017 | 85 | 75 | 0.26 | 0.0020 | 0.44 |
|  |  |  |  | 2020 | 29 | 24 | 0.24 | 0.0018 |  |
|  |  |  |  | 2021 | 6 | 6 |  |  |  |
|  | San Joaquin Basin | Sharktooth Creek (SHK) | 3.2 | 2021 | 40 | 37 | 0.22 | 0.0020 | 0.42 |
|  |  | Stairway Creek (SWC) | 3.5 | 2021 | 38 | 32 | 0.18 | 0.0019 | 0.39 |

<sup>1</sup> Occupied/available habitat for within-basin streams are derived from Ryan and Nicola (1976) and as reported in FWS (2004).
<sup>2</sup> Occupied/available habitat for out-of-basin streams are from FWS (2004).

Fin clips were taken from live fish and were dried on Whatman qualitative filter paper and placed in coin envelopes. We extracted DNA from fin clips with a magnetic bead-based protocol (Ali et al. 2016) and quantified using Quant-iT PicoGreen dsDNA Reagent (Thermo Fisher Scientific) with an FLx800 Fluorescence Reader (BioTek Instruments). We prepared RAD libraries for 476 samples (Supplementary materials S1) using the *Sbf1* restriction enzyme following the BestRAD protocol in Ali et al. (2016), and then pooled and sequenced samples on an Illumina NovaSeq 6000 at the DNA Technologies and Expression Analysis Core at the University of California, Davis with paired-end 150-bp reads. We used fastq-multx 1.4.2 to demultiplex the sequencing data with an exact match with plate barcodes (https://github.com/brwnj/fastq-multx) using BarcodeSplitListBestRadPairedEnd.pl frmo Ali et al. (2016). Demultiplexed data were aligned to the Rainbow Trout reference genome (NCBI Bioproject PRJNA623027) using the bwa-mem algorithm in the Burrows-Wheeler Aligner 0.7.17 (Li and Durbin 2009). We used SAMtools 1.7.2 (Li et al. 2009) to sort, remove duplicates, and count the mapped reads for each individual.

Individuals with low sequencing success were determined by both coverage in terms of mapped reads and the ability to call genotypes. Genotypes were called by running ANGSD 1.9 (Korneliussen et al. 2014) with the SAMtools 1.7.2 genotype likelihood model (-GL 1) and a uniform prior (-doPost 2) to identify genotypes with mapping scores above 30 (-minMapQ 30) and base quality scores above 20 (-minQ 20). Individuals with <600,000 mapped reads and <16,000 called SNPs were omitted from downstream analyses.

### SNP genotyping and filtering

We re-called genotypes for all individuals that passed screening described above using ANGSD 1.9 (Korneliussen et al. 2014) with the SAMtools model (-GL 1), inferring major and minor alleles directly from genotype likelihoods (-doMajorMinor 1), estimating posterior genotype probability assuming a uniform prior (-doPost 2) and estimating allele frequencies assuming a fixed major allele and minor allele (-doMaf 1). We only used reads with a mapping score ≥30 (-minMapQ 30) and bases with a quality score ≥20 (-minQ 20). Furthermore, we only included SNPs with a minor allele frequency of at least 0.01 (-minMaf 0.01) that were represented in at least 60% of samples. To reduce linkage disequilibrium, we used BCFtools 1.14 (Li et al. 2009) with default parameters to prune SNPs that had Lewontin’s D > 0.1.

An initial principal component analysis (PCA) using PCAngsd (Meisner and Albrechtsen 2018) revealed a strong signal associated with the RAD library from which samples originated instead of the biological patterns expected based on previous work in this system (Finger et al. 2013). We reduced this batch effect by identifying loci shared across libraries, excluding loci with high *F_ST_* values between libraries, and excluding loci with unusually low or high sequencing depth (see Supplementary Materials for details).

Approximately 88 million years ago a whole-genome duplication occurred in salmonid fish, leaving a large proportion of paralogous loci in descendant forms such as cutthroat trout species (Macqueen and Johnston 2014). Given the lineage-specific aspect of rediploidization apparent in salmonids, we identified paralogous loci specific to PCT instead of relying on homology with known residually tetrasomic genomic regions in other taxa (e.g., Robertson et al. 2017; Blumstein et al. 2020). We used HDplot to remove paralogous SNPs called in each population separately to avoid filtering out SNPs associated with genetic population structure (McKinney et al. 2017). We retained SNPs with a heterozygosity <0.45 and balanced allele read ratios (Li 2011; Danecek et al. 2021). Hereafter, we refer to the set of SNPs retained after the filtering steps described thus far as our ‘**RAD-seq SNPs**’.

Finally, to select candidate loci for a genetic monitoring SNP panel that were present and polymorphic in all sampled populations, we further filtered the RAD-seq SNPs to only retain loci with a mean allele ratio (D) between 0.4 and 0.6 for individuals in every population (McKinney et al. 2017). This set of SNPs is referred to as our ‘**panel SNPs**’.

### in silico SNP panel validation

To validate the ability of our panel SNPs to replicate patterns of genetic structure and diversity observed using the full RAD-seq dataset, we compared principal component analysis (PCA) results and individual heterozygosity results produced using each set of SNPs. We verified that the panel SNPs and RAD-seq SNPs produced similar individual heterozygosity results by performing a Wilcoxon signed rank test using the R package rstatix with a strict Bonferroni adjusted-P value (Kassambara 2020). Detailed methods for performing the PCA and calculating individual heterozygosity can be found in the next two sections.

### Range-wide genetic structure

To investigate the range-wide population genetic structure of the refuge populations, we conducted a PCA with all the samples that passed filtering. For our PCA, we generated a covariance matrix using PCAngsd (Meisner and Albrechtsen 2018), then used the eig() function in R 4.1.2 to compute the eigen matrix and visualized the PCA with ggplot2 (Wickham 2016). We further characterized genetic structure by performing an admixture analysis using NGSadmix (Skotte et al. 2013). We ran 10 iterations of each K value (the number of genetic groups) from 1 to 10, then we used the Evanno method (Evanno et al. 2005) to select optimal K and we visualized admixture proportions for the optimal K value and its adjacent K values.

To quantify the degree of genetic differentiation among refuge populations, we estimated pairwise *F_ST_* values. We used ANGSD 1.9 to generate a VCF file (-dovcf 1) with genotypes for all panel SNPs, and then used VCFtools 0.1.14 (Danecek et al. 2011) to calculate the Weir and Cockerham *F_ST_* for each SNP locus. We considered missing and negative *F_ST_* values as 0 and calculated the mean *F_ST_* value among loci in each population. To examine differentiation within populations over time, we also calculated pairwise *F_ST_* between different years for the same location for two locations, upper Silver King Creek and North Fork Cottonwood Creek. Pairwise *F_ST_* values were visualized using the R package ggplot2 (Wickham 2016).

### Range-wide genetic diversity

We calculated the mean and variance of genome-wide heterozygosity for each individual, using ANGSD (Korneliussen et al. 2014) to compute a folded site frequency spectrum (SFS) based on genotype likelihoods for the panel SNPs. We then estimated heterozygosity from the folded SFS as the proportion of sites with intermediate allele frequencies. We performed a Wilcoxon signed rank test with a Bonferroni adjustment in the R package rstatix (Kassambara 2020) to detect significant differences in individual heterozygosity among the refuge populations and between years for the two locations sampled multiple times. After detecting a significant difference in individual heterozygosity between sampling years for North Fork Cottonwood Creek, we downsampled 2017 to match the sample size for 2020/2021 and redid Wilcoxon signed rank testing to ensure the difference was not an artifact of variation in sample size.

### Detecting spawning after a translocation

In 2017, an ongoing drought in North Fork Cottonwood Creek reduced streamflow to dangerous levels. In response, managers conducted an emergency translocation, removing (and sampling) 88 adult PCT from North Fork Cottonwood Creek and releasing them in upper Silver King Creek (USFWS 2020). Fin clip samples were collected in upper Silver King Creek two years and four years after translocation, from one-year-old fish (60 mm or less; Titus and Calder 2009) and mixed-age fish, respectively, to discover any changes resulting from the influx of translocated fish. This event enabled us to explore how our markers may be used after a translocation event by evaluating (1) whether our SNP panel can reliably detect offspring admixed from two populations, and (2) whether our SNP panel can identify direct descendants of translocated fish.

To detect North Fork Cottonwood Creek ancestry in upper Silver King Creek fish collected after the translocation event, we performed admixture analysis restricted to the donor and recipient populations: North Fork Cottonwood Creek in 2017, and upper Silver King Creek in 2019 and 2021. We used the SNP panel to generate a beagle file in ANGSD and used NGSadmix to generate the q-value matrix for the selected populations when K = 2.

To identify direct descendants of fish translocated from North Fork Cottonwood Creek to upper Silver King Creek, we conducted parentage analysis with the program COLONY2 (Jones and Wang 2010) using SNP panel genotypes reformatted with the snpR function write_colony_input() (Hemstrom and Jones 2023). COLONY2 requires genotypes from potential parents and offspring to conduct parentage analysis. For potential parents, we used all translocated North Fork Cottonwood Creek individuals that passed sequence quality filters (n = 75). For potential offspring in this analysis, we used genotypes of all individuals collected in upper Silver King Creek in 2021 (n = 87) and any individuals identified as admixed collected in upper Silver King Creek in 2019 (n = 4). For negative controls (to ensure that COLONY was not selecting spurious parents), we included as potential offspring two individuals collected from Coyote Valley Creek in 2019. We set the probability of parents being present in our dataset to 0.5, used the full likelihood method, and set genotype error rate to 0.1 and dropout rate to 0.01. We assumed outbreeding and selected random, polygamous mating for the mating system.

As sampling of all possible parents from either population was not exhaustive, we also applied NewHybrids (Anderson and Thompson 2002) to identify potential descendants of translocated fish from North Fork Cottonwood Creek to upper Silver King Creek. NewHybrids can identify F1s, F2s, or backcrossed individuals. To do so, we selected 10 fish to represent North Fork Cottonwood Creek based on being identified as a parent in Colony or having a high North Fork Cottonwood Creek admixture proportion (*Q* ∼ 1.00). We applied the same reasoning to identify 10 fish representative of the upper Silver King Creek genetic background. With these 20 representative fish of North Fork Cottonwood Creek and upper Silver King Creek, we conducted an analysis with NewHybrids of the fish identified as admixed individuals and, therefore, likely descendants of translocated North Fork Cottonwood Creek individuals using the gl.nhybrids function of dartR (Gruber et al. 2018). The panel SNP genotype calls were read into R with the read.vcfR function of the vcfR package and converted to a genlight object with vcfR2genlight (Knaus et al. 2017). NewHybrids has a limit of 200 markers so we applied filtering to the dataset to reduce the number of markers and enrich the dataset with markers most informative for hybrid inference. We filtered for a MAF of 0.15 across North Fork Cottonwood Creek and upper Silver King Creek with the gl.filter.maf and gl.filter.na functions of dartR, then extracted the representative and potential descendant fish. We required individuals to have genotypes for >70% of the retained loci to be included in the NewHybrids analysis. NewHybrids itself was driven by the gl.nhybrids function of dartR, designating representative fish from North Fork Cottonwood Creek as ‘Parent population 0’ (P0) and representative fish from upper Silver King Creek as ‘Parent population 1’ (P1). A 300,000 step burn-in and 50,000 steps were specified.

## Results

### Sequencing and SNP filtering

After filtering individuals for sequencing and mapping quality, 402 out of 476 individuals were kept for further analysis (Table 1, Supplementary materials Table S1). We used 212,592 SNPs in the initial range-wide PCA, where we observed a strong batch effect (Supplementary materials Figure S1). After filtering for shared loci, *F_ST_* outliers, and sequencing depth, our dataset had 29,390 SNPs, and a PCA showed the batch effect was substantially reduced (Supplementary materials Figure S1). Due to small sample sizes for Corral Valley Creek 2019 and North Fork Cottonwood Creek 2021, we combined Corral Valley Creek 2019 and Corral Valley Creek 2020, as well as North Fork Cottonwood Creek 2020 and 2021, each into a single population for all analyses, labeled as Corral Valley Creek 2019 and North Fork Cottonwood Creek 2020, respectively (Table 1). After merging all the non-paralogous SNPs identified by HDplot in each population, we selected 12,085 SNPs for the RAD-seq SNP dataset and 1,716 SNPs for the candidate panel SNPs dataset. After pruning for linkage disequilibrium, our final RAD-seq SNP dataset included 6,187 loci, and our candidate panel SNPs dataset included 1,114 loci.

### In silico SNP panel validation

The PCAs generated by RAD-seq SNPs (Figure 2A, B) and panel SNPs (Figure 2C, D) were similar, but the panel SNPs produced a clearer resolution of the expected population structure based on Finger et al. (2013) and the panel SNPs had a higher proportion of variance explained by the first three PC axes (10% versus <5%). For most of our populations, we found no significant difference in mean individual heterozygosity estimates from panel SNPs compared to RAD-seq SNPs (Figure 2E); however, we observed slightly more variance in panel SNPs (Figure 2C, Supplementary materials Table S3). An exception was Stairway Creek samples from 2021, which showed a modest but statistically significant difference in mean individual heterozygosity between datasets (0.18 vs. 0.21; ∼15% higher in the panel SNP dataset).

**Figure 2.**
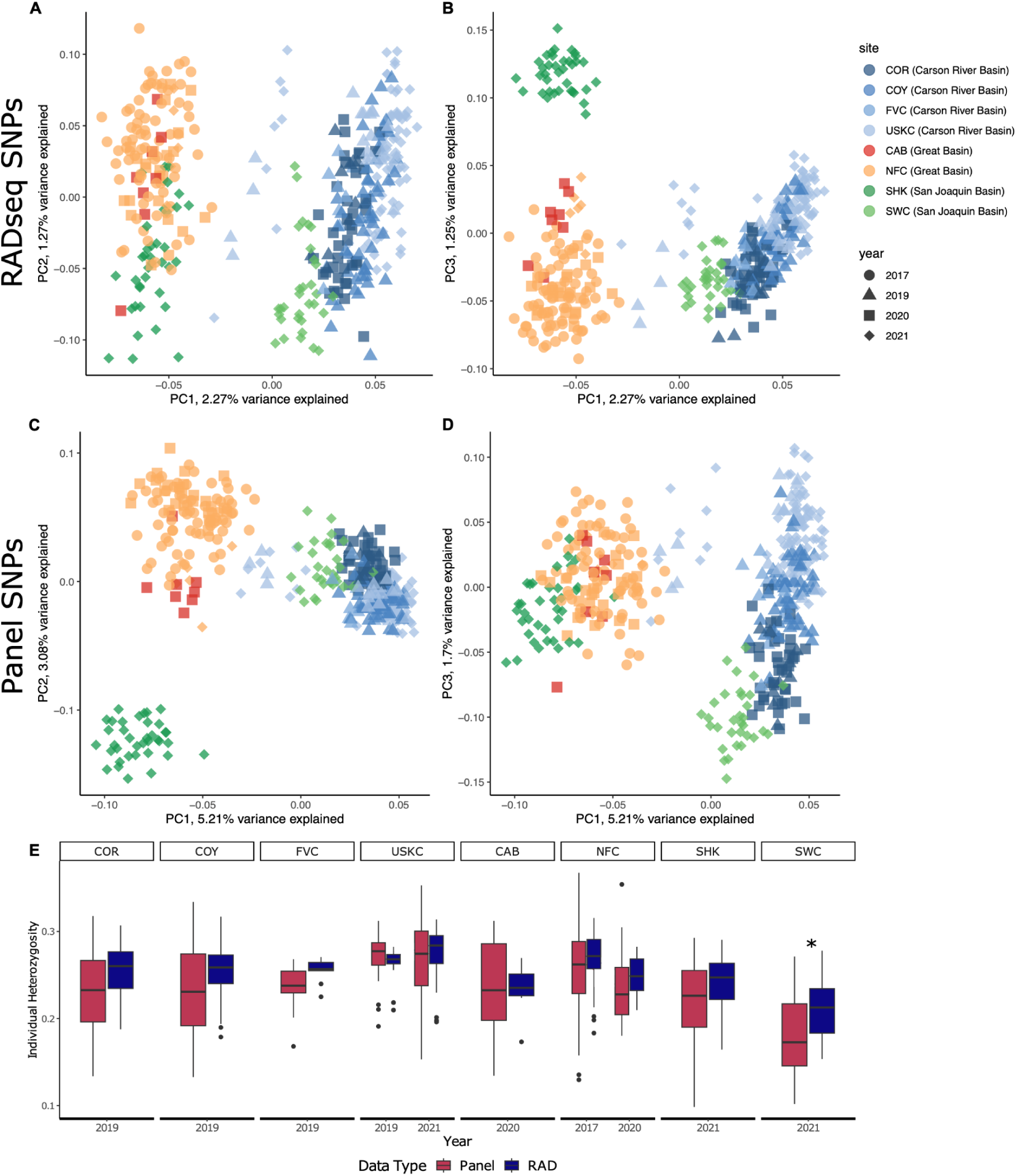
*In silico* validation of our candidate panel SNPs. Principal component analysis (PCA) based on 6,187 unlinked, non-paralogous RADseq SNPs are shown for PC1 vs. PC2 (A) and PC1 vs. PC3 (B). Comparable PCA plots based on 1,114 unlinked, non-paralogous panel SNPs are shown for PC1 vs. PC2 (C) and PC1 vs. PC3 (D). Panel (E) compares individual heterozygosity estimates from RADseq and panel SNPs; significant differences are indicated by an asterisk. Full names for creeks can be found in Table 1.

### Range-wide genetic structure

The first three PC axes in our PCA explained 10% of the genetic variation in the SNP panel dataset (Figure 3A, B). In PC1 and PC2, the four out-of-basin populations are separated from each other as well as from the within-basin populations, with Stairway Creek clustering the closest to within-basin populations. Cabin Creek and North Fork Cottonwood Creek, both from the Great Basin, are more closely related to each other than to the geographically distant Sharktooth Creek. Within-basin populations are more closely related to each other than out-of-basin populations. PC3 reveals more genetic differentiation among the within-basin populations (Figure 3B). Corral Valley Creek is more unique from the other within-basin populations, while Fly Valley Creek, Coyote Valley Creek, and upper Silver King Creek are clustered more closely together. Twelve individuals from upper Silver King Creek sampled in 2019 or 2021 were located intermediately between within-basin and out-of-basin populations.

**Figure 3.**
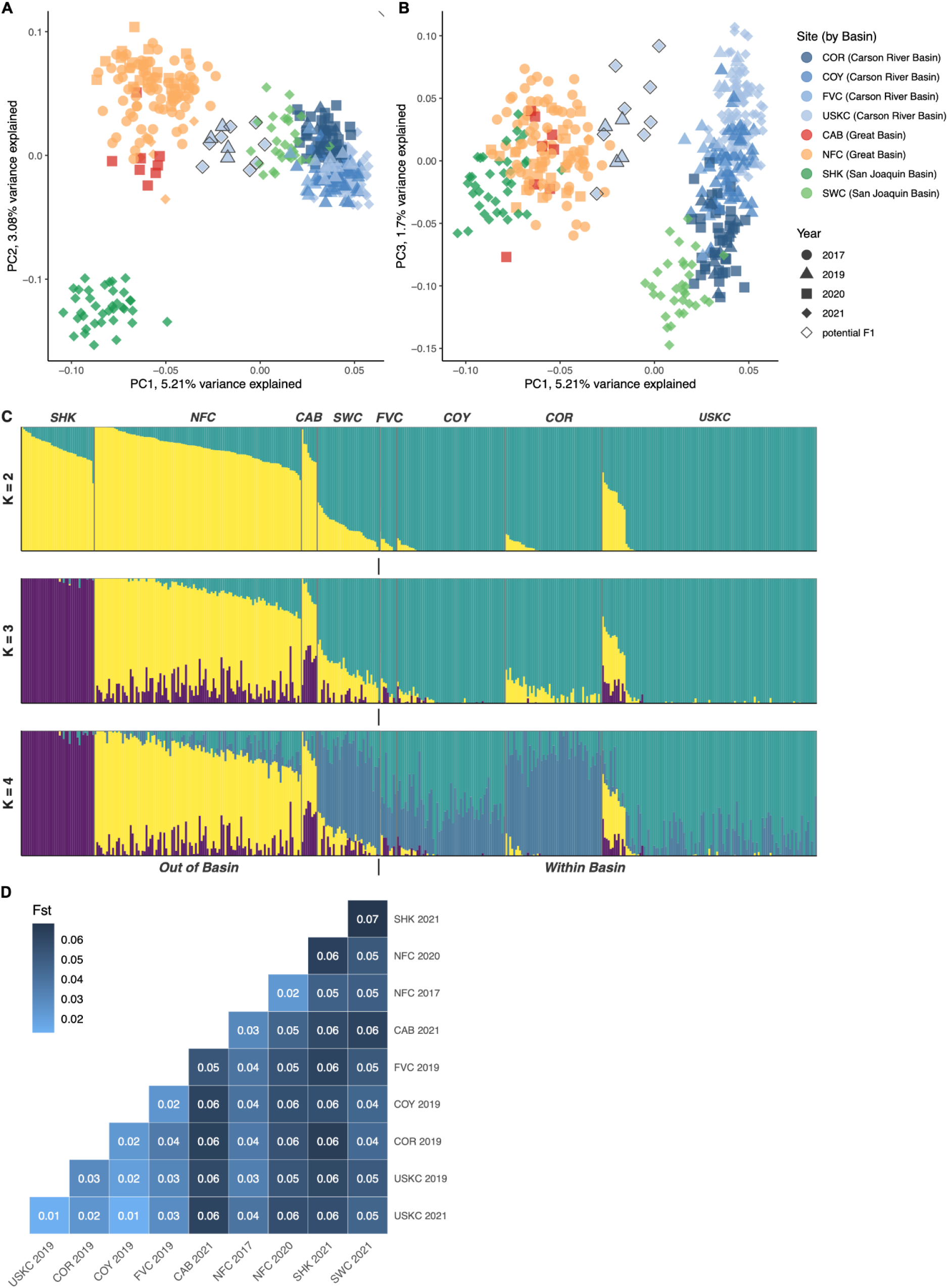
Genetic structure analyses of Paiute Cutthroat Trout using 1,114 panel SNPs, including a principal component analysis: PC1 vs. PC2 (A) and PC1 vs. PC3 (B), an admixture analysis for K=2–4 (C), and pairwise *F_ST_* values among refuge populations and among sampling years within the same populations (D). Full names for creeks can be found in Table 1. Shapes with black outlines in the PCAs are fish with NGSadmix-inferred ancestry assigning to both North Fork Cottonwood Creek and upper Silver King Creek (i.e., potential F1 offspring after translocation), which we further evaluated in Figure 5.

Our admixture analysis showed the highest mean log-likelihood at K = 1 (Supplementary materials Figure S4). We visualized the admixture results for K = 2–4 to examine whether any biologically meaningful patterns of admixture emerged at higher K values, even though K = 1 was statistically favored. At K = 2, three of four out-of-basin populations (Sharktooth Creek, North Fork Cottonwood Creek, Cabin Creek) were assigned largely to one cluster, while all four within-basin populations were assigned largely to the other cluster, and out-of-basin Stairway Creek showed more admixture between the two clusters (Figure 3C). Most of the upper Silver King Creek samples were partitioned into the same cluster as the other within-basin populations, but 12 upper Silver King Creek individuals were admixed with both clusters. At K = 3, Sharktooth Creek formed a distinct cluster (Figure 3C). At K = 4, Corral Valley Creek separated from the other within-basin populations, and Stairway Creek, Fly Valley Creek, and Coyote Valley Creek also had a high proportion of admixed ancestry in this cluster (Figure 3C). These results largely align with our PCA results.

*F_ST_* values were generally low among all refuge populations (range: 0.01–0.07, Figure 3D). Similar to the PCA and admixture analysis, Sharktooth Creek had the highest *F_ST_* values when compared with all other populations. Upper Silver King Creek in 2021 had the lowest *F_ST_* compared to all other populations. *F_ST_* values within populations sampled at multiple time points were lower than *F_ST_* values among populations: *F_ST_* between upper Silver King Creek in 2019 and 2021 was 0.01, and *F_ST_* between North Fork Cottonwood Creek in 2017 and 2020 was 0.02.

### Range-wide genetic diversity

Mean individual heterozygosity in each refuge population ranged from 0.18 to 0.27 (Figure 4, Table 1). Stairway Creek had significantly lower individual heterozygosity than all other populations except for Cabin Creek (Supplementary materials Table S4). In contrast, upper Silver King Creek 2021 had significantly higher heterozygosity than all other populations. Sharktooth Creek was also significantly different from six populations, with individuals generally exhibiting elevated heterozygosity compared to other populations. There was no significant difference in heterozygosity in upper Silver King Creek between 2019 and 2021, indicating temporal stability in genetic diversity. North Fork Cottonwood Creek, however, showed a significant decline in heterozygosity between 2017 and 2020 samples, even after downsampling 2017 to the same sample size as 2020.

**Figure 4.**
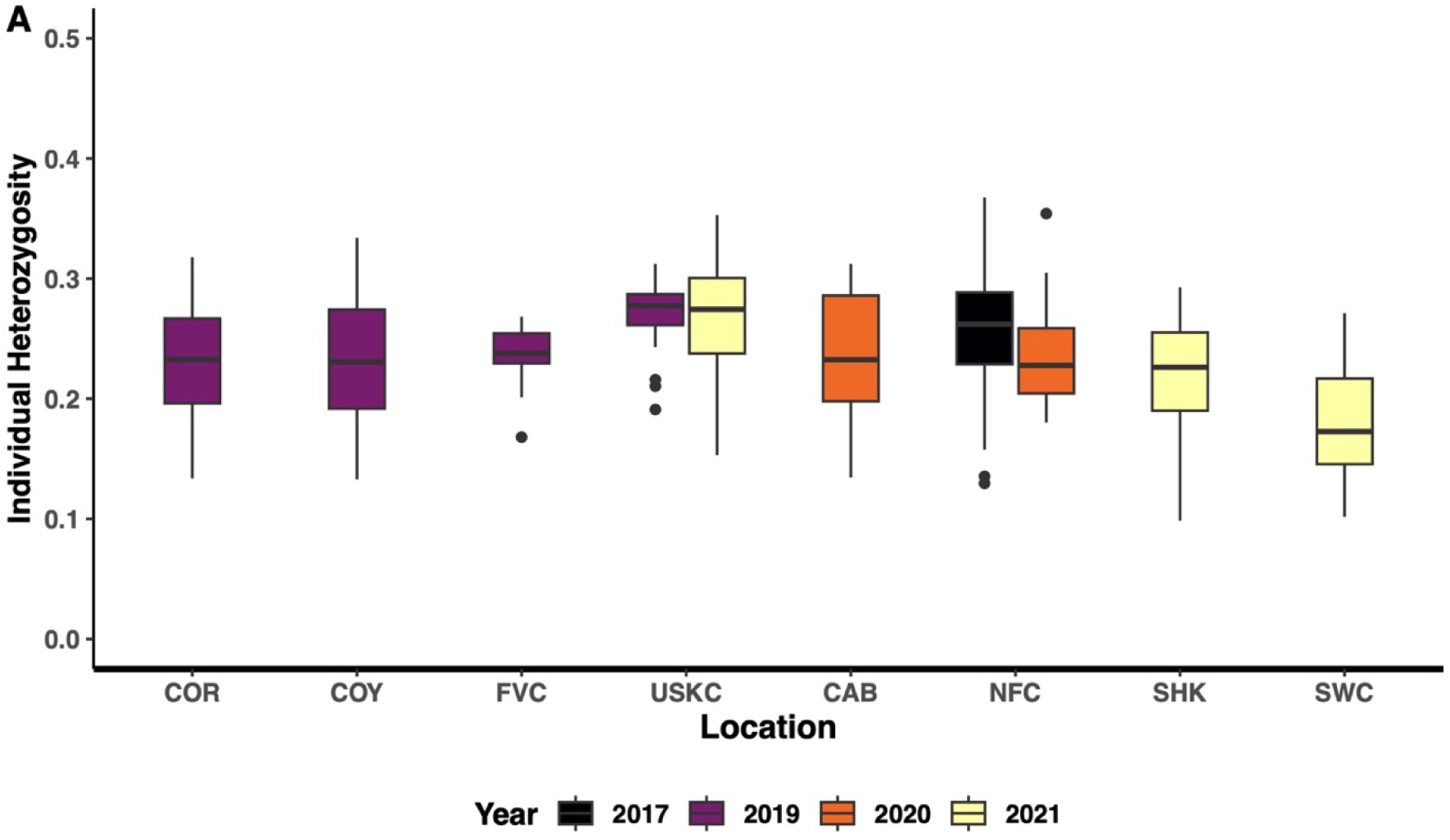
Individual heterozygosity estimates based on genotype data from panel SNPs for Paiute Cutthroat Trout samples in each refuge population, colored by sampling year. Mean and variance values, as well as full names for creeks, can be found in Table 1.

### Translocation monitoring

Our admixture analysis of fish sampled in North Fork Cottonwood Creek and upper Silver King Creek revealed 12 upper Silver King Creek fish with more than 38% ancestry from the North Fork Cottonwood Creek donor population, while all others had <10% (Figure 5A). Four of these mixed-ancestry individuals were collected in 2019 and the other eight were collected in 2021.

**Figure 5.**
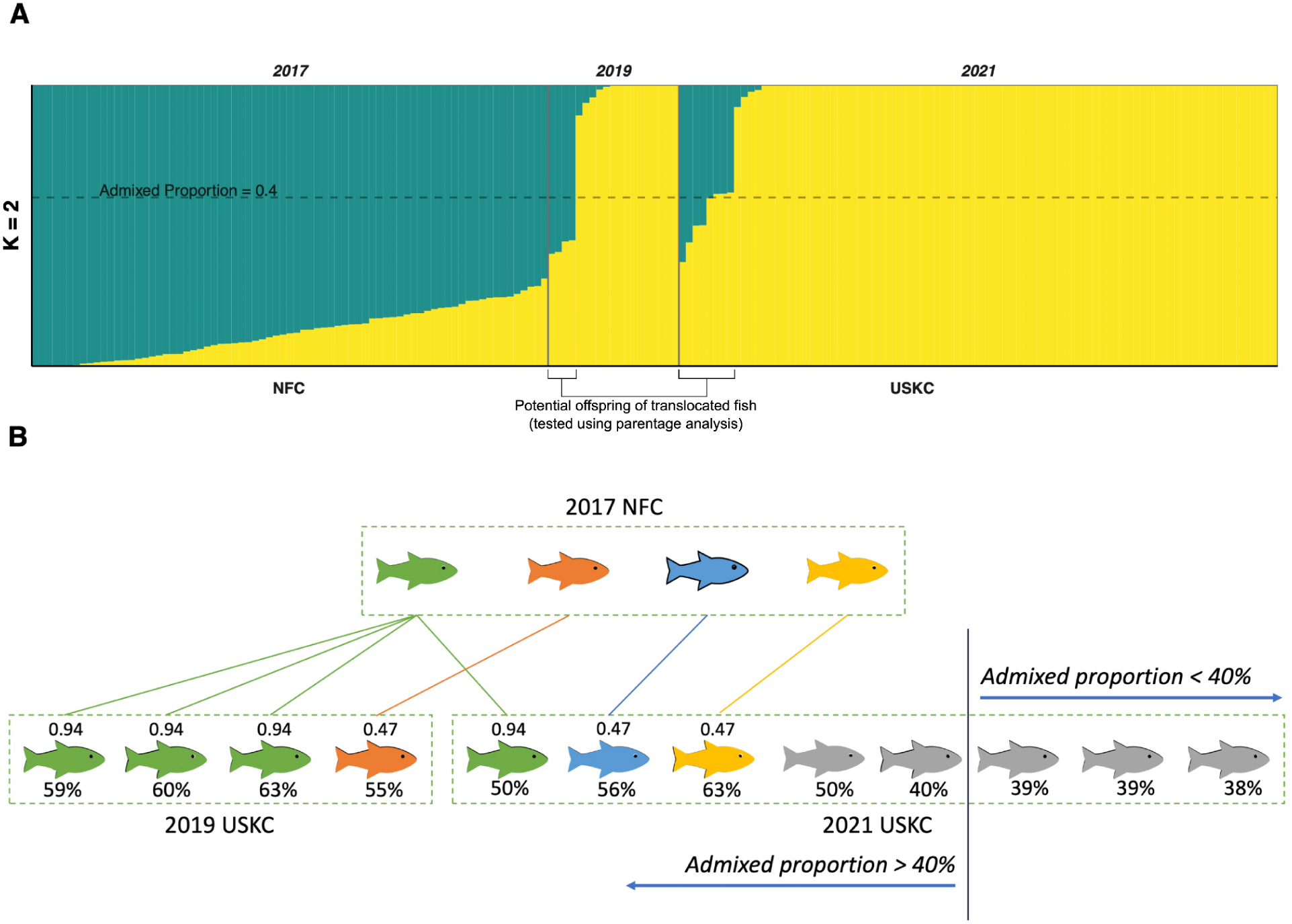
Ancestry of Paiute Cutthroat Trout sampled in upper Silver King Creek (USKC) after the 2017 translocation of 88 fish from North Fork Cottonwood Creek (NFC) to USKC, using 1,114 panel SNPs, based on parentage analysis performed using the software program COLONY. (A) Admixture proportions when K=2 of translocated fish and fish sampled in USKC after translocation. (B) Parent-offspring relationships among translocated fish and USKC fish with NFC ancestry. Each fish symbol represents an individual. Fish of the same color and connected by lines have an inferred parent-offspring relationship. The posterior probability of each parentage assignment is indicated above the fish and the percentage of NFC ancestry is indicated below the fish. Fish in grey denote fish with no parent identified.

After running parentage analysis for these 12 fish, we successfully identified a North Fork Cottonwood Creek-origin parent for seven of them (Figure 5B). Three out of four 2019 individuals and one 2021 individual were identified as having the same translocated parent with a 95% probability. One 2019 individual and two other 2021 individuals each assigned to one translocated parent, though the probability of each pairing was only 47%. The three individuals with 38%–39% North Fork Cottonwood Creek ancestry failed to have any parents identified. Our negative control samples from an external population did not assign to any parents.

After filtering the panel SNPs for analysis with NewHybrids, we used 177 SNPs to analyze 32 individuals. This included 10 pure North Fork Cottonwood Creek fish and 10 pure upper Silver King Creek fish for reference, plus the 12 fish that our admixture analysis identified as having mixed ancestry. All 20 reference fish retained assignments to their respective populations, and all 12 admixed fish were indicated to be F1 offspring (Supplemental Figure S5). Of the 12 admixed fish, the seven fish for which Colony identified parents had a mean probability of assignment to the F1 class of 0.92 and median value of 0.96. The other five admixed fish for which Colony could not identify parents had a mean probability of assignment to the F1 class of 0.99 and median value of 0.99.

## Discussion

In this study, we applied RAD sequencing to generate a genome-wide SNP dataset for PCT. From these, we identified a subset of 1,114 SNPs that can be developed into a GT-seq style amplicon sequencing panel for long-term, range-wide genetic monitoring of PCT. The panel SNPs recapitulated the findings of the RADseq SNPs and offered some improvements, particularly in differentiating closely related, low diversity populations. Importantly, we established a genetic baseline for future genetic monitoring. The SNP panel also identified offspring of individuals translocated between populations of PCT and detected reduced genetic diversity in the donor population after the translocation event, demonstrating the effectiveness of the candidate panel for monitoring and species management.

### Range-wide genetic structure

The patterns of population genetic structure we observed align with the complex translocation history outlined in stocking records from the California Department of Fish and Wildlife, which are summarized by USFWS (2004). *F_ST_* values were low among all populations, with the lowest differentiation among within-basin populations (Figure 3). Similarities among the four within-basin populations can be explained by shared source populations and reciprocal translocations. For example, Fly Valley Creek was established with fish from Corral Valley Creek and Coyote Valley Creek in 1947. Subsequently, in the 1970s and 1990s, Corral and Coyote Valley creeks and upper Silver King Creek were chemically treated to remove hybridized trout after which Fly Valley Creek served as a source population for re-establishment. This culminated in the genetic similarity among all four within-basin populations. Compared to the within-basin populations, out-of-basin populations have had comparatively fewer translocations. The North Fork Cottonwood Creek population was originally established from upper Silver King, Corral Valley, and Coyote Valley creeks in 1946. North Fork Cottonwood Creek was then utilized as a source population to establish two other out-of-basin populations, Cabin Creek and Sharktooth Creek, both in 1968. Sharktooth Creek was stocked not only from North Fork Cottonwood Creek but also from the now-extirpated out-of-basin Delaney Creek, which was itself established from the now-extirpated within-basin Four Mile Canyon Creek. Our PCA shows a high degree of overlap between North Fork Cottonwood Creek and Cabin Creek, reflecting their shared origin, whereas Sharktooth Creek is more genetically distinct from both. In contrast, Stairway Creek was established in 1972 with fish from Delaney Creek, and shows a closer genetic relationship to within-basin populations. This suggests that while Sharktooth Creek has diverged from both its source populations and others, Stairway Creek retains stronger genetic ties to the within-basin lineage. Our SNP panel effectively captures the genetic patterns shaped by historic translocations, providing a baseline for measuring population structure in PCT.

The broad scale patterns of genetic structure were comparable between the panel SNPs and RADseq SNPs, demonstrating that the signal from panel SNPs successfully captures the signal from RADseq SNPs. The panel SNPs further excelled in revealing a gradient of differentiation among within-basin populations, whereas the RADseq PCA showed a more muddled relationship among them (Figure 2A-D). Despite employing many fewer loci, the panel SNPs matched or outperformed the RADseq SNPs in distinguishing among PCT refuge populations.

### Range-wide genetic diversity

After comparing genetic diversity metrics from our panel SNPs to those from the full set of 6,187 RAD-seq SNPs, we found that the panel produced comparable estimates of genetic diversity at the population level for each refuge population (Table 1, Figure 2). Patterns of genetic diversity inferred from mean individual heterozygosity using our panel SNPs were broadly consistent with those reported in Finger et al. (2013), which used microsatellites to calculate expected heterozygosity. For example, Stairway Creek had the lowest levels of genetic diversity across both marker types (0.18 in this study; 0.39 in Finger et al. 2013), while populations such as upper Silver King Creek and North Fork Cottonwood Creek exhibited comparatively higher diversity in both datasets (0.27 and 0.26 in this study; 0.53 and 0.44 in Finger et al. 2013). Upper Silver King Creek and North Fork Cottonwood Creek appear to have received the largest number of founder individuals based on the best available records (CDFW stocking records as summarized in USFWS 2004), which may explain the higher heterozygosity values. Aside from the low mean heterozygosity observed in Stairway Creek, all of the other creeks had relatively similar levels of heterozygosity, regardless of whether they were within- or out-of-basin. This is particularly interesting given the variation in occupied stream lengths (and assumed related variation in census sizes) among creeks (Table 1).

### Translocation monitoring

Translocation is a widely used conservation strategy aimed at bolstering small, isolated populations, mitigating inbreeding, or maintaining genetic diversity across a species’ range. For species like the PCT, which persists solely in a network of small, genetically depauperate refuge populations, translocations are a tool for recovery and possibly a necessary intervention to preserve evolutionary potential and prevent local extirpation. Using the panel SNPs, we identified fish with a mix of upper Silver King Creek and North Fork Cottonwood Creek ancestry, suggesting that at least some North Fork Cottonwood Creek-origin fish successfully spawned after being translocated into upper Silver King Creek in 2017. The 12 admixed fish that we found had 38%–59% North Fork Cottonwood Creek ancestry based on our NGSadmix results, and our NewHybrids analysis confirmed that these 12 fish are highly likely to be F1 offspring from one translocated North Fork Cottonwood Creek parent and one upper Silver King Creek parent. Our Colony parentage analysis went a step further by identifying a total of four North Fork Cottonwood Creek-origin parents for seven of the 12 F1 offspring. The other five F1 fish failed to have North Fork Cottonwood Creek parents identified. One potential reason is that their North Fork Cottonwood Creek parent is one of the 11 individuals translocated from North Fork Cottonwood Creek that did not pass our filtering threshold for inclusion in our analyses. Overall, these findings confirm successful spawning after a translocation effort and demonstrate the utility of the SNP dataset for detecting such an event.

Translocations can affect the donor populations demographically and genetically due to the removal of individuals. In this study, our sample consisted of drought-affected fish that were removed from North Fork Cottonwood Creek and placed into upper Silver King Creek. Census size estimates for North Fork Cottonwood Creek in the past have ranged from 576 to 704 (Wong 1975; UFWS 2004), so, conservatively, 88 fish removed from a population of 704 is 12.5%. We found that genetic diversity in North Fork Cottonwood Creek decreased from 0.26 in 2017 to 0.24 in 2020/2021, even after accounting for sample size differences. It is possible that the removal of 88 fish plus drought conditions in the intervening years contributed to this decrease. Under scenarios such as assisted gene flow, translocations seek to have minimum impact on the donor population. In this case, however, North Fork Cottonwood Creek was deemed to have water flows so low that the population was at risk of extirpation or a severe bottleneck. Furthermore, moving out-of-basin fish to the Silver King Creek watershed is part of a longer-term plan to restore lower Silver King Creek and repatriate PCT to their native habitat.

Overall, translocations may be an effective management strategy to maintain the genetic diversity of PCT. Our detection of interpopulation F1 offspring after the translocation of fish from North Fork Cottonwood Creek to upper Silver King Creek demonstrates initial evidence that translocated fish will spawn with recipient population fish. This is salient in light of significant management actions taken since sample collection for this study. For example, three additional translocations occurred to repopulate PCT into its historical range in lower Silver King Creek after nonnative trout removal: in 2019, 30 PCT were translocated from Coyote Valley Creek; in 2020, 44 PCT were translocated from Corral Valley Creek; and in 2021, an additional 52 PCT were translocated from upper Silver King Creek (USFWS 2025). Genetic monitoring will assess the value and success of these translocations and guide managers on how effective colonizing lower Silver King Creek with fish from multiple locations is with regards to population viability and boosting genetic diversity.

Westslope Cutthroat Trout (*Oncorhynchus lewisi*) serves as an example of a well-studied system with a successful history of translocations. The species currently exists in widespread fragmented habitats, and, as a result, populations may have significant structure, and some have low genetic variation (Bell et al. 2025). Kovach et al. (2022) confirmed that translocations successfully increased genetic variation in recipient populations. However, because PCT are not highly structured and diversity levels are similar among extant populations, the benefit of assisted gene flow may be less dramatic than what was observed in Westslope Cutthroat Trout.

## Conclusion

Recovery of PCT is active and ongoing, at this time relying heavily on monitoring and control of nonnative trout species. Ultimately, the goal is to have a resilient, sustaining population in lower Silver King Creek and the continued existence of refuge populations as a safeguard against extinction. Lower Silver King Creek continues to face a multitude of threats including interspecific hybridization with Rainbow Trout, small population sizes, and effects of climate change and its restoration and on-going management will be a multi-year, if not perpetual, effort. Conservation managers often have limited time and resources; the genetic baseline results herein and periodic monitoring will inform effectiveness of past and future management decisions, enumerating changes over time to genetic diversity and divergence among refuge populations.

## Supporting information

All supplement files

## Acknowledgements

We thank the California Department of Fish and Wildlife, United States Fish and Wildlife Service, and United States Forest Service personnel for collecting the genetic samples. We thank Sean O’Rourke and Mary E. Badger for help with extracting DNA and building RAD sequencing libraries. The sequencing was carried out by the DNA Technologies and Expression Analysis Core at the UC Davis Genome Center, supported by NIH Shared Instrumentation Grant 1S10OD010786-01. Funding provided by CDFW grant #P1920001.

## Data availability

Genetic data and associated metadata are deposited in the SRA (BioProject PRJNA1353977; https://www.ncbi.nlm.nih.gov/sra).

## Ethics statement

Tissue samples for this project were obtained from individuals listed in the acknowledgements. Collections for this study followed institutional guidelines and permitting.

## Supplement (separate google doc)

Table S1 – sample metadata [upload separately from the rest of the supplement]

Table S2 – batch effect sample sizes by library

Table S3 – pops with sample sizes and RADseq vs. panel mean ind het

Table S4 – pairwise comparison of pop mean individual heterozygosity values

Fig S1 – batch effect PCAs

Fig S2 – batch effect Fst along genome

Fig S3 – batch effect read depth and Fst

Fig S4 – delta N values for admixture analysis

Fig S5 – NewHybrids plot

