## Supplementary material for "Use of SNPs in a low diversity system for genetic monitoring and identifying a successful translocation event in the Paiute Cutthroat Trout (*O. henshawi seleniris*)": All supplement files

### I. Identifying and reducing the sequencing batch effect

#### Methods

*Identifying a sequencing batch effect*

We performed an initial principal component analysis (PCA) as described in our paper, and found that the PCA results did not reflect expected population genetic structure based on Finger et al. (2013). Rather, the PCA revealed a strong signal associated with the RAD library from which samples originated, especially for samples from Library 9 (Figure S1-A, D). According to Finger et al. (2013), within-basin populations have more similar genetic variation than out-of-basin populations, so we created two subsets of our data that each included samples in Library 9 and the other libraries that were sourced from the same populations in a similar year (Table S2). The first dataset included samples from all within-basin populations that were located both in Library 9 and other libraries (Library 1 and 6). The second dataset included individuals from one out-of-basin population, North Fork Cottonwood Creek, which were in Library 9 and other libraries (Library 7 and 8). This allowed us to identify variants associated with library batch differences rather than a true biological signal. We called genotypes for within-basin populations and North Fork Cottonwood Creek separately using the same parameters listed in the main text using ANGSD 1.9 (Korneliussen et al. 2014). We only called genotypes with data for 60% of individuals and generated vcf files (-dovcf 1) for each subset.

*SNP filtering to reduce the batch effect*

To address the sequencing batch effect, we filtered our SNP datasets based on three criteria: loci shared across libraries, $F_{ST}$ at loci among libraries, and sequencing depth.

When we discovered SNP loci for each library independently, some SNPs only appeared in one or several libraries instead of all of them. To assess if these non-shared SNPs contributed to the observed batch effect, we used bedtools (Quinlan and Hall 2010) to identify SNPs in common between Library 9 (i.e., the plate with the strongest batch effect) and the other libraries for North Fork Cottonwood Creek and within-basin populations separately. We combined all the intersected SNPs that were shared between North Fork Cottonwood Creek dataset and within-basin populations dataset, and then called genotypes for all range-wide samples at this combined SNP list, then performed PCA to assess if the batch effect was reduced.

To further investigate the cause of the batch effect, we calculated pairwise $F_{ST}$ and sequencing depth for each locus in the samples that were influenced by the batch effect. We identified $F_{ST}$ outliers associated with sequencing libraries for the North Fork Cottonwood Creek and within-basin populations datasets. We used VCFtools 0.1.14 (Danecek et al. 2011) to compare Weir and Cockerham $F_{ST}$ for every locus in our combined SNP list. We identified SNPs with $F_{ST}$ values >1.5x the interquartile range as outliers (Grubbs 1969). We then extracted sequencing depth for each SNP for each individual using VCFR 1.12.0 (Knaus and Grünwald 2017), calculated the mean depth of each SNP, and conducted a Wilcoxon signed rank test in R package rstatix 0.7.0 (Kassambara 2020) to compare mean depth of $F_{ST}$ outliers and non-outliers. To reduce the batch effect, we removed all SNPs that we identified as $F_{ST}$ outliers or had a mean depth <5 or >50.

### Results

We used 212,592 SNPs in the initial range-wide PCA, where we observed a strong batch effect (Figure S1-A, D). Filtering our SNP dataset to only include loci in common between Library 9 and the other libraries resulted in a dataset with 95,932 SNPs, but a PCA using this dataset revealed little improvement on the original batch effect (Figure S1-B, E). We identified 6,533 out of 54,049 SNPs as $F_{ST}$ outliers in North Fork Cottonwood Creek, and 6,680 out of 48,711 SNPs $F_{ST}$ outliers in within-basin populations. These $F_{ST}$ outliers had significantly lower depth than the non-outlier SNPs in both North Fork Cottonwood Creek and within-basin populations (Figure S2). After filtering for $F_{ST}$ outliers and depth of coverage, our dataset included 29,390 SNPs and a PCA showed the batch effect was substantially reduced (Figure S1-C, F). We proceeded with further filtering and analysis using this dataset, as described in the main text.

### Discussion

In this study, we encountered a strong artificial effect of library preparation in our RAD sequencing data, which interfered with data analysis to identify informative SNPs for genetic monitoring. In populations with low levels of genetic diversity or little genetic differentiation among populations, batch effects may present a stronger signal than the biological pattern, making it difficult to discern true genomic relationships among populations. Batch effects can arise in a variety of ways that are not mutually exclusive, including preparation of sequencing libraries by different personnel, errors in standardizing input DNA concentrations, or stochasticity in PCR steps during the process of next generation sequencing (De-Kayne et al. 2021; Leigh et al. 2018; O'Leary et al. 2018). Previous studies have demonstrated how batch effect can be caused by poly-G tails, base quality score miscalibration, coverage differences, reference bias and alignment errors caused by difference in read type and read length, and DNA degradation (De-Kayne et al. 2021; Lou and Therkildsen 2022; O'Leary et al. 2018). Removing $F_{ST}$ outliers associated with different batches (RAD sequencing libraries, in our case) can sufficiently remove the batch effect for populations with minimal population structure that are sequenced across multiple libraries. This method may be especially valuable for filtering SNPs when samples from the same populations are partitioned into multiple libraries. If no samples in the library with a batch effect originate from the same location as the samples in the other libraries, performing genotype calling in one batch and applying the SNPs on another batch can also be an effective way to address the issue (Lou and Therkildsen 2022).

Table S2. The number of samples from each population and sampling year included in each RAD-seq library. Populations highlighted in red represent samples influenced by the batch effect, and populations highlighted in yellow denote samples not influenced by the batch effect that were used as the comparison group for filtering out loci based on outlier FST values. Note: Some plates had additional samples not listed here that were excluded from final analyses; no excluded samples were relevant for analyzing the batch effect.

| **Library** | **Population** | **Sampling Year** | **Sample size** | **Sample size after filter** |
|---|---|---|---|---|
| Plate 1 | Upper Silver King Creek | 2021 | 96 | 87 |
| Plate 2 | Upper Silver King Creek | 2021 | 20 | 0 |
| Plate 6 | Coyote Valley Creek | 2019 | 34 | 32 |
| | Corral Valley Creek | 2020 | 44 | 39 |
| | Sharktooth Creek | 2021 | 9 | 8 |
| Plate 7 | Sharktooth Creek | 2021 | 31 | 29 |
| | Stairway Creek | 2021 | 38 | 32 |
| | North Fork Cottonwood Creek | 2020 | 19 | 14 |
| | North Fork Cottonwood Creek | 2021 | 6 | 6 |
| Plate 8 | North Fork Cottonwood Creek | 2020 | 10 | 10 |
| | Cabin Creek | 2020 | 8 | 8 |
| Plate 9 | North Fork Cottonwood Creek | 2017 | 48 | 47 |
| | Coyote Valley Creek | 2019 | 15 | 15 |
| | Corral Valley Creek | 2019 | 7 | 7 |
| | Upper Silver King Creek | 2019 | 18 | 17 |
| | Fly Valley Creek | 2019 | 5 | 5 |
| Plate 10 | Coyote Valley Creek | 2019 | 12 | 8 |
| | Corral Valley Creek | 2019 | 8 | 3 |
| | Upper Silver King Creek | 2019 | 6 | 5 |
| | North Fork Cottonwood Creek | 2017 | 37 | 28 |
| | Fly Valley Creek | 2019 | 5 | 4 |
| *Total* | | | *476* | *404* |

Table S3. Refuge population sampling information and individual heterozygosity values. The number of SNPs discovered in each population for RAD-seq and panel SNPs before and after filtering paralogous SNPs by HDplot, and the mean and variance of individual heterozygosity in each population and year generated with 6,197 RAD-seq SNPs and 1,114 panel SNPs. North Fork Cottonwood Creek 2020 and 2021, and Corral Valley Creek 2019 and 2020 were each merged into one population (labeled as North Fork Cottonwood Creek 2020 and Corral Valley Creek 2019 in the analyses) because of the low sampling size in North Fork Cottonwood Creek 2021 and Corral Valley Creek 2019.

| Type | Basin | Population | Sample collection year | Sample size | Selected SNPs | | Mean heterozygosity | | Variance heterozygosity | |
|---|---|---|---|---|---|---|---|---|---|---|
| | | | | | RAD-seq SNPs | Panel SNPs | RAD-seq SNPs | Panel SNPs | RAD-seq SNPs | Panel SNPs |
| Within-basin | Carson River Basin | Corral Valley Creek (COR) | 2019 | 10 | 8,423 | 305 | 0.26 | 0.24 | 0.0005 | 0.002 |
| | | | 2020 | 39 | | | 0.26 | 0.23 | 0.0009 | 0.002 |
| | | Coyote Valley Creek (COY) | 2019 | 55 | | | 0.26 | 0.23 | 0.0010 | 0.003 |
| | | Fly Valley Creek (FVC) | 2019 | 9 | | | 0.26 | 0.23 | 0.0002 | 0.001 |
| | | Upper Silver King Creek (USKC) | 2019 | 20 | | | 0.26 | 0.27 | 0.0004 | 0.0001 |
| | | | 2021 | 87 | | | 0.27 | 0.27 | 0.0007 | 0.002 |
| Out-of-basin | Great Basin | Cabin Creek (CAB) | 2020 | 8 | 6,823 | 662 | 0.23 | 0.24 | 0.0008 | 0.004 |
| | | North fork Cottonwood Creek (NFC) | 2017 | 75 | 8,054 | 621 | 0.27 | 0.26 | 0.0007 | 0.002 |
| | | | 2020 | 24 | | | 0.25 | 0.23 | 0.0004 | 0.002 |
| | San Joaquin Basin | Sharktooth Creek (SHK) | 2021 | 37 | 6,898 | 589 | 0.22 | 0.24 | 0.0011 | 0.002 |
| | | Stairway Creek (SWC) | 2021 | 32 | 5,383 | 365 | **0.18** | **0.21** | 0.0012 | 0.002 |

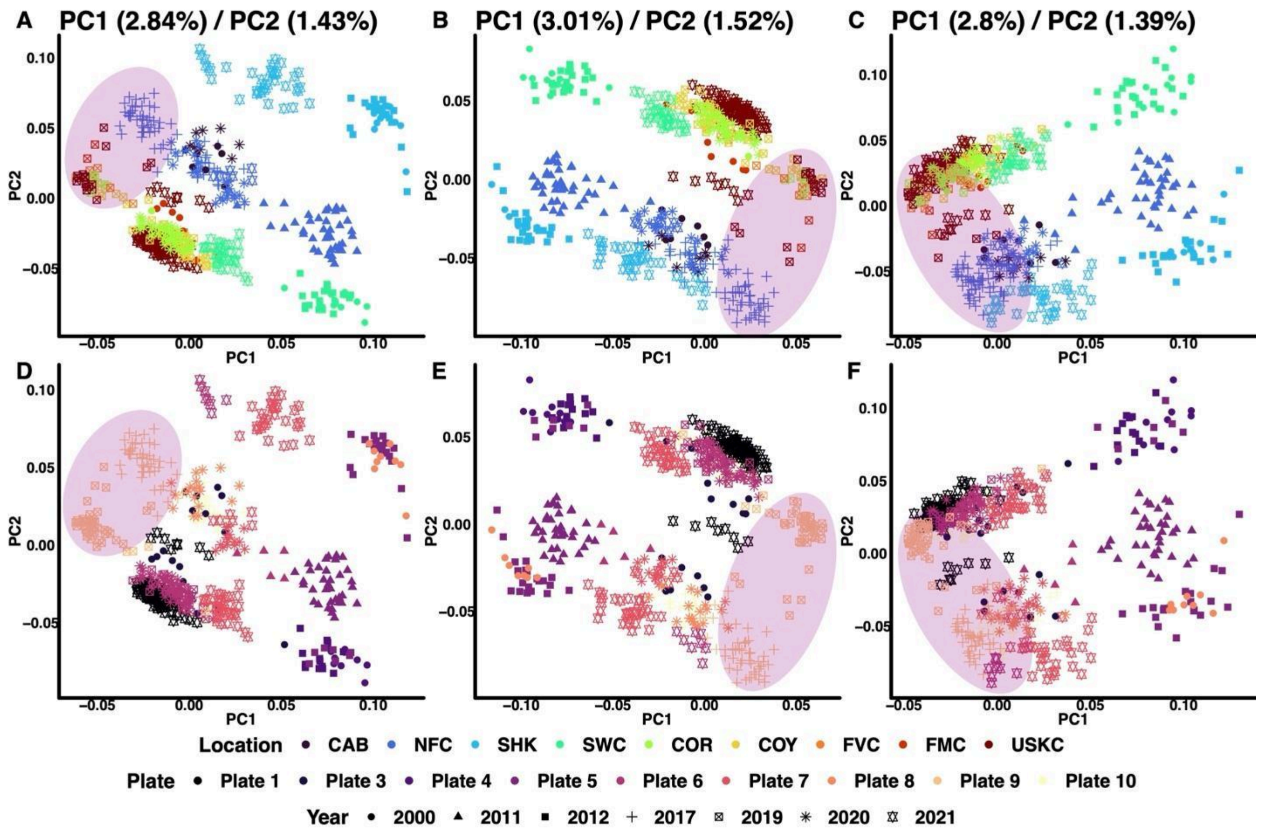


Figure S1. PCAs with sample colors representing population or sequencing library and shape representing sampling year. PCAs colored by population or sequencing library were generated using (A, D) SNPs with no filter, (B, E) after removing SNPs not shared across libraries, and (C, F) after removing FST outliers and SNPs with unusually high or low sequencing depth. Individuals most impacted by the batch effect are highlighted with pink shaded ovals. Note: Samples collected from 2000–2012 are shown here for reference but were excluded from final analyses; this exclusion is reflected throughout the main text.

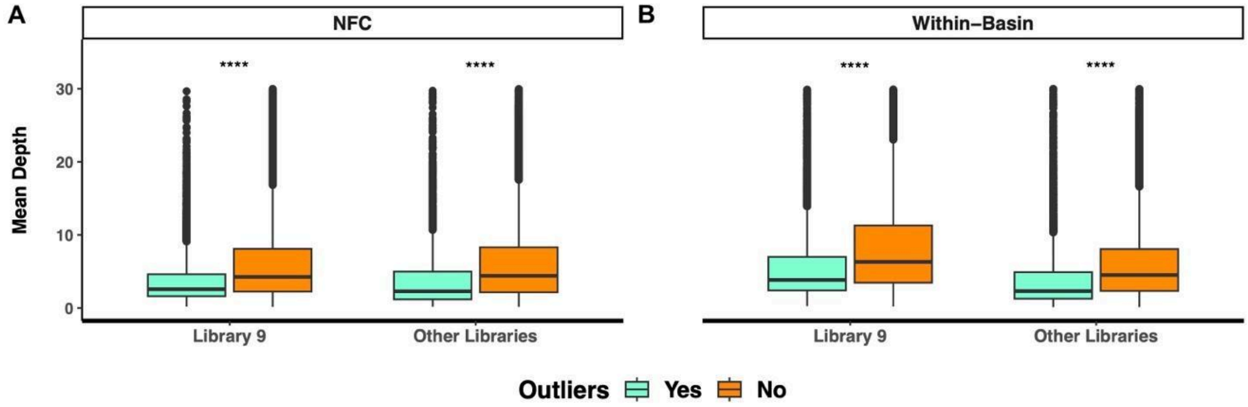


Figure S2. Mean sequencing depth in $F_{ST}$ outliers and non-outlier SNPs between Library 9 and other libraries for (A) north fork Cottonwood Creek (NFC) and (B) all within-basin populations (P-values: ****=0.0001).

II. Other supplementary materials

Table S4. The pairwise comparison of individual heterozygosity values in each pair of populations each year. Sample size 1 corresponds to population 1, and sample size 2 corresponds to population 2. P-value significance values: ns: p > 0.05, *: p <= 0.05, **: p <= 0.01, ***: p <= 0.001, ****: p <= 0.0001.

| Population 1 | Population 2 | Sample size 1 | Sample size 2 | p-value | Adjusted p-value | Significance of adjusted p-value |
|---|---|---|---|---|---|---|
| CAB 2020 | COR 2019 | 16 | 98 | 0.367 | 1 | ns |
| CAB 2020 | COY 2019 | 16 | 110 | 0.397 | 1 | ns |
| CAB 2020 | FVC 2019 | 16 | 18 | 0.251 | 1 | ns |
| CAB 2020 | NFC 2017 | 16 | 150 | 0.006 | 0.27 | ns |
| CAB 2020 | NFC 2020 | 16 | 48 | 0.673 | 1 | ns |
| CAB 2020 | SHK 2021 | 16 | 74 | 0.748 | 1 | ns |
| CAB 2020 | SWC 2021 | 16 | 64 | 0.002 | 0.09 | ns |
| CAB 2020 | USKC 2019 | 16 | 38 | 0.008 | 0.36 | ns |
| CAB 2020 | USKC 2021 | 16 | 174 | 0.001 | 0.045 | * |
| COR 2019 | COY 2019 | 98 | 110 | 0.999 | 1 | ns |
| COR 2019 | FVC 2019 | 98 | 18 | 0.763 | 1 | ns |
| COR 2019 | NFC 2017 | 98 | 150 | 0.000139 | 0.006 | ** |
| COR 2019 | NFC 2020 | 98 | 48 | 0.345 | 1 | ns |
| COR 2019 | SHK 2021 | 98 | 74 | 0.02 | 0.9 | ns |
| COR 2019 | SWC 2021 | 98 | 64 | 9.36E-11 | 4.21E-09 | **** |
| COR 2019 | USKC 2019 | 98 | 38 | 0.002 | 0.09 | ns |
| COR 2019 | USKC 2021 | 98 | 174 | 3.50E-08 | 1.58E-06 | **** |
| COY 2019 | FVC 2019 | 110 | 18 | 0.813 | 1 | ns |
| COY 2019 | NFC 2017 | 110 | 150 | 0.000176 | 0.008 | ** |
| COY 2019 | NFC 2020 | 110 | 48 | 0.424 | 1 | ns |
| COY 2019 | SHK 2021 | 110 | 74 | 0.025 | 1 | ns |
| COY 2019 | SWC 2021 | 110 | 64 | 1.01E-10 | 4.55E-09 | **** |
| COY 2019 | USKC 2019 | 110 | 38 | 0.002 | 0.09 | ns |
| COY 2019 | USKC 2021 | 110 | 174 | 9.13E-08 | 4.11E-06 | **** |
| FVC 2019 | NFC 2017 | 18 | 150 | 0.007 | 0.315 | ns |
| FVC 2019 | NFC 2020 | 18 | 48 | 0.592 | 1 | ns |
| FVC 2019 | SHK 2021 | 18 | 74 | 0.179 | 1 | ns |
| FVC 2019 | SWC 2021 | 18 | 64 | 4.73E-06 | 0.0002 | *** |
| FVC 2019 | USKC 2019 | 18 | 38 | 0.000212 | 0.010 | ** |
| FVC 2019 | USKC 2021 | 18 | 174 | 0.000489 | 0.022 | * |
| NFC 2017 | NFC 2020 | 150 | 48 | 3.89E-05 | 0.002 | ** |
| NFC 2017 | SHK 2021 | 150 | 74 | 4.68E-09 | 2.11E-07 | **** |
| NFC 2017 | SWC 2021 | 150 | 64 | 1.12E-19 | 5.04E-18 | **** |

| | | | | | | |
|---|---|---|---|---|---|---|
| NFC 2017 | USKC 2019 | 150 | 38 | 0.685 | 1 | ns |
| NFC 2017 | USKC 2021 | 150 | 174 | 0.036 | 1 | ns |
| NFC 2020 | SHK 2021 | 48 | 74 | 0.281 | 1 | ns |
| NFC 2020 | SWC 2021 | 48 | 64 | 3.92E-08 | 1.76E-06 | **** |
| NFC 2020 | USKC 2019 | 48 | 38 | 5.08E-05 | 0.002 | ** |
| NFC 2020 | USKC 2021 | 48 | 174 | 1.69E-07 | 7.61E-06 | **** |
| SHK 2021 | SWC 2021 | 74 | 64 | 7.59E-06 | 0.0003 | *** |
| SHK 2021 | USKC 2019 | 74 | 38 | 8.68E-07 | 3.91E-05 | **** |
| SHK 2021 | USKC 2021 | 74 | 174 | 1.53E-12 | 6.89E-11 | **** |
| SWC 2021 | USKC 2019 | 64 | 38 | 5.37E-13 | 2.42E-11 | **** |
| SWC 2021 | USKC 2021 | 64 | 174 | 9.16E-22 | 4.12E-20 | **** |
| USKC 2019 | USKC 2021 | 38 | 174 | 0.095 | 1 | ns |

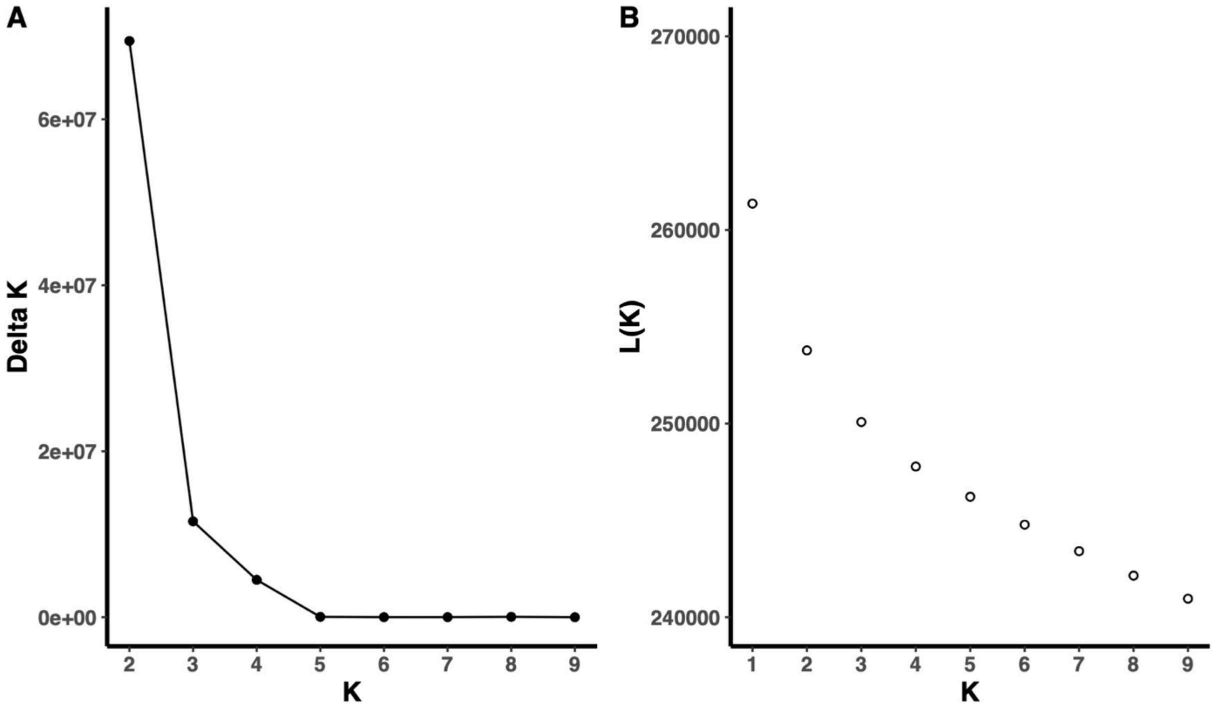


Figure S4. Optimal K selected by Evanno method for the range-wide admixture analysis using 1,114 panel SNPs. (A) Delta K was calculated from K=2–9. (B) Abstract value of mean log likelihood of each K calculated from K=1–9.

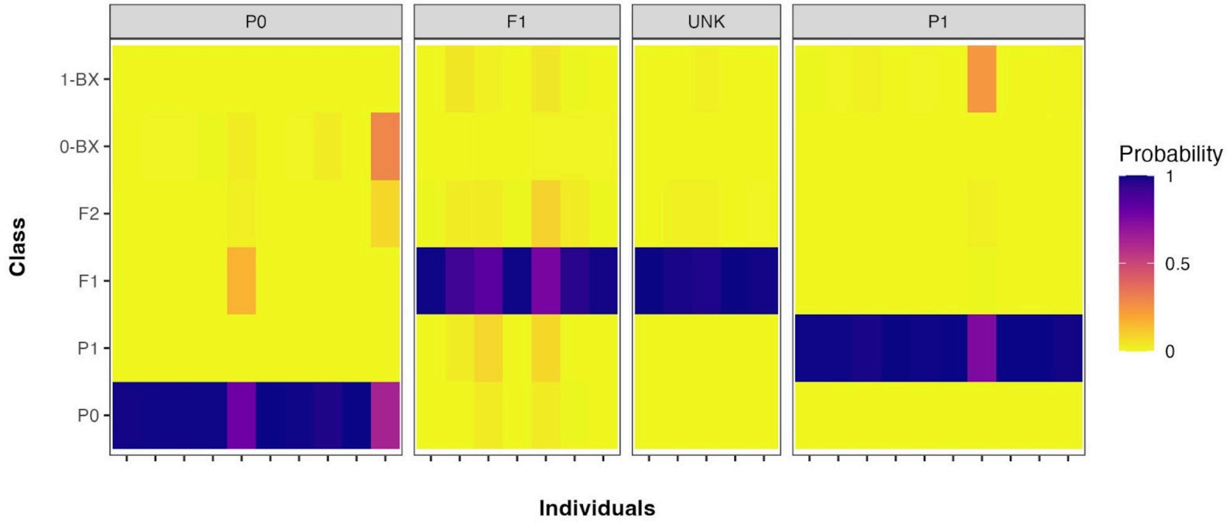


Figure S5. NewHybrids analysis of potential offspring of North Fork Cottonwood Creek translocated individuals. Individuals are plotted along the *x*-axis and split into categories. P0 is North Fork Cottonwood Creek background, P1 is upper Silver King Creek background, F1 are fish identified by Colony to be offspring of translocated individuals and UNK are fish supported by admixture analyses to be possible descendants of translocated North Fork Cottonwood Creek fish.
